# Bacteriophages antagonize cGAS-like immunity in bacteria

**DOI:** 10.1101/2022.03.30.486325

**Authors:** Erin Huiting, Januka Athukoralage, Jingwen Guan, Sukrit Silas, Héloïse Carion, Joseph Bondy-Denomy

## Abstract

The recently discovered <u>c</u>yclic-oligonucleotide-<u>b</u>ased <u>a</u>nti-phage <u>s</u>ignaling <u>s</u>ystem (CBASS) is related to eukaryotic cGAS-STING anti-viral immunity and is present in diverse prokaryotes. However, our understanding of how CBASS detects, inhibits, and co-evolves with phages is limited because CBASS function has only been studied in reconstituted heterologous systems. Here, we identify a phage-encoded CBASS antagonist (*acbIIA1,* <u>a</u>nti-<u>cb</u>ass type <u>II-A</u> gene <u>1</u>) necessary for phage replication in the presence of endogenous CBASS immunity in *Pseudomonas aeruginosa*. *acbIIA1* homologs are encoded by numerous lytic and temperate phages infecting Gram-negative bacteria. Deletion of *acbIIA1* renders multiple phages susceptible to CBASS, but phages can then escape immune function via mutations in the major capsid gene. These mutants suggest that CBASS is activated by, or targets, the late-expressed phage capsid. Together, we establish a native model system to study CBASS and identify a common phage-encoded CBASS antagonist, demonstrating that CBASS is a bona fide anti-phage immune system in nature.

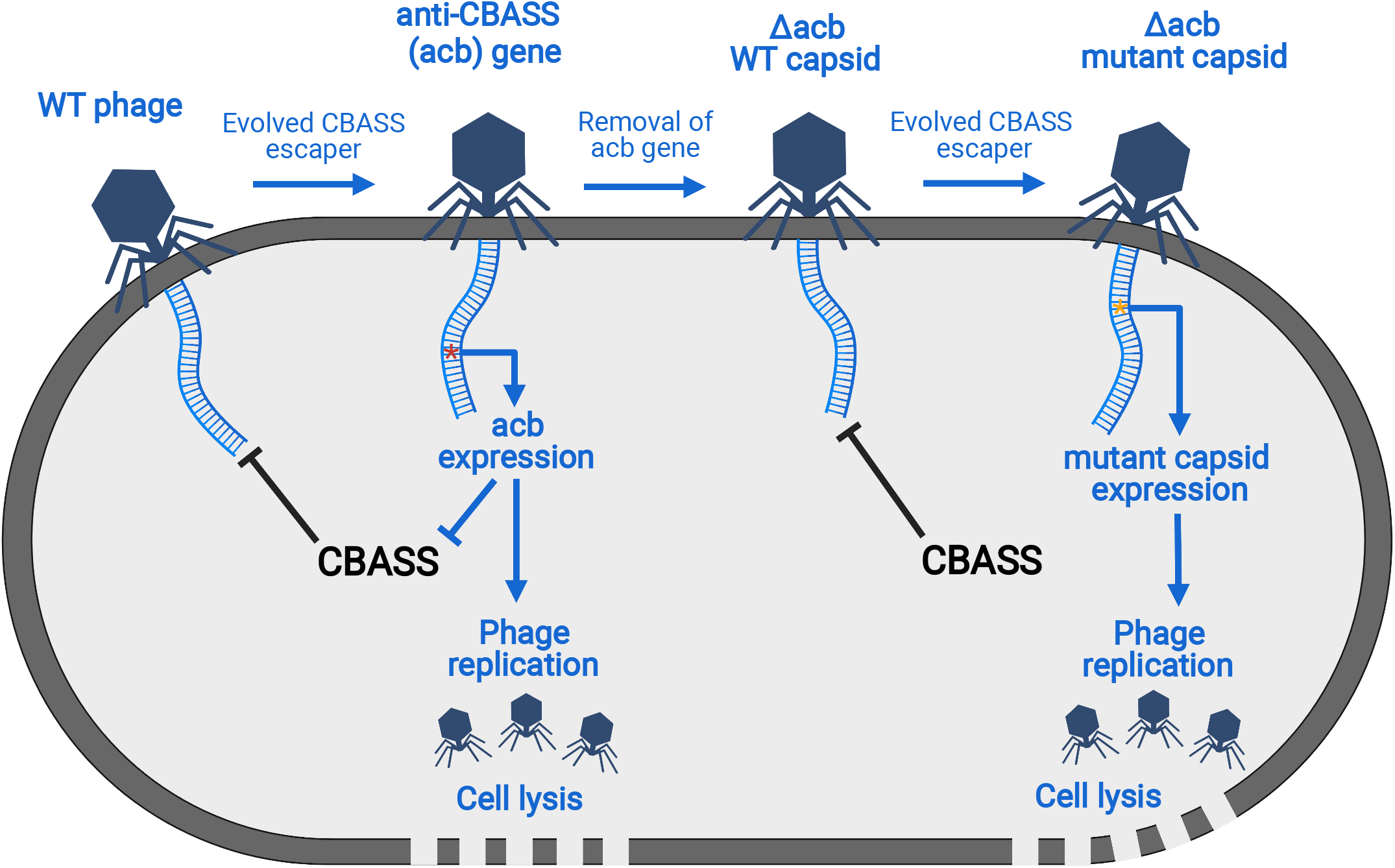
Graphical Abstract.

## Introduction

Sensing specific macromolecules produced or possessed by viruses is a fundamental tenet of anti-viral immune systems across all kingdoms of life (Barbalat et al., 2011; Margolis et al., 2017). In mammalian cells, viral double-stranded DNA (dsDNA) is directly bound by cyclic GMP-AMP synthase (cGAS) in the cytoplasm (Li et al., 2013; Wu et al., 2013). The activated cGAS enzyme then produces cyclic GMP-AMP (cGAMP) signaling molecules that bind to the adaptor protein stimulator of interferon genes (STING) and induces a type I interferon response (Ishikawa et al., 2008; Burdette et al., 2011). Recently, thousands of cGAS-like enzymes were identified across the entire bacterial domain and biochemically characterized (Whiteley et al., 2019), drawing back the evolutionary origin of cGAS-based immune systems by approximately 2 billion years (Margulis 1995). These enzymes, called <u>c</u>GAS/<u>D</u>ncV-like <u>n</u>ucleotidyltransfer<u>ases</u> (CD-NTases) produce cyclic oligonucleotides, such as cGAMP, during bacteriophage (phage) infection. When assayed in heterologous expression systems, cyclic oligonucleotides activate a downstream effector (Cohen et al., 2019; Lau et al., 2020; Ye et al., 2020; Lowey et al., 2020; Duncan-Lowey et al., 2021), which is proposed to kill the cell and prevent phage replication (Lopatina et al., 2020). This strategy of bacterial immunity was coined <u>c</u>yclic-oligonucleotide-<u>b</u>ased <u>a</u>nti-phage <u>s</u>ignaling <u>s</u>ystems (CBASS) (Cohen et al., 2019). CD-NTases and effectors are encoded adjacent to each other and are recognized as the ‘core’ genes (Cohen et al., 2019) present in all CBASS types (Millman et al., 2020). Type I CBASS operons encode just the CD-NTase and effector pair, while Type II and III operons encode additional ‘signature’ <u>C</u>D-NTase-<u>a</u>ssociated <u>p</u>roteins (Cap) (Lowey et al., 2020; Millman et al., 2020). Type II signature proteins Cap2 and Cap3 are of unknown function whereas the Type III signature Cap proteins possess CBASS regulatory functions (Burroughs et al., 2015; Lau et al., 2020; Ye et al., 2020). Despite this level of understanding of CBASS, how phages induce cyclic-oligonucleotide production and whether CBASS is a bona fide immune system that places selective pressure on phages *in situ* is unknown.

The identification of bacterial strains that endogenously use CBASS to protect against phage infection will aid in our understanding of CBASS function(s), mechanisms, and regulation. *Pseudomonas aeruginosa* is a generalist microbe that survives in many niches, has a diverse phage population, and is an important human pathogen that resists many antimicrobial drugs. Expanding our understanding of *P. aeruginosa* phage-host interactions is critical to developing new antimicrobial therapeutics. Previous analyses (Millman et al., 2020), coupled with our own bioinformatics, reveal that 252 distinct *P. aeruginosa* strains encode >300 CBASS operons spanning Type I-III systems, which encode one of four different effectors (phospholipase (A), transmembrane (B), endonuclease (C), phosphorylase (E)) and at least four different cyclic-oligonucleotides (cGAMP, cyclic tri-AMP, cyclic UMP-AMP, cyclic UMP-GMP; Whiteley et al., 2019). This suggests that CBASS is important for *P. aeruginosa* fitness and that it may be a good model organism for studying phage-CBASS interactions. Here, we identified a naturally active Type II-A CBASS system (cGAMP producing) with a phospholipase effector in *P. aeruginosa* that limits phage replication. We next identified a widespread phage gene that is necessary for phage replication in the presence of CBASS. Lastly, we isolated phages that escape CBASS with mutations in their major capsid gene. This work provides the first direct evidence of phage-CBASS co-evolution, demonstrating a robust arms race between the two.

## Results

### Endogenous anti-phage CBASS function in *Pseudomonas aeruginosa*

To identify strains with naturally functional CBASS immunity, CBASS operons were deleted from four *P. aeruginosa* strains (possessing representatives of three common CBASS types: Type II-A, II-C, and III-C) and the mutants were screened against a diverse panel of 70 *P. aeruginosa*-specific phages spanning 23 different genomic families and four different morphologies (Myoviridae, Siphoviridae, Podoviridae, and Inoviridae) (Ha and Denver, 2018). In one strain (BWHPSA011), the deletion of the Type II-A CBASS locus (ΔCBASS, Fig. 1A) resulted in >4 orders of magnitude increase in titer of the dsDNA podophage PaMx41 (Figure 1B-C). Phage protection was restored when all four CBASS genes [CapV (phospholipase effector), CdnA (cyclase), Cap2 (E1/E2), and Cap3 (JAB)] were complemented on a plasmid (Figure 1D). PaMx41 shares >96% nucleotide identity with phages PaMx33, PaMx35, and PaMx43, which exhibited resistance to CBASS (Figure 1B-1C, S1). However, episomal overexpression of the CBASS operon reduced their titer by 1-2 orders of magnitude (Figure 1D). To determine which genes are necessary for CBASS anti-phage activity, we generated chromosomal mutants of each CBASS gene that is predicted to disrupt catalytic activity (Cohen et al., 2019): (i) CapV S48A, (ii) CdnA D87A/D89A, (iii) Cap2 C450A/C453A, and (iv) Cap3 E38A. The mutations in CapV, CdnA and Cap2 abolished anti-phage activity, while the Cap3 mutation did not (Figure 1E, S1). These data demonstrate that *P. aeruginosa* BWHPSA011 (herein Pa011) CBASS-based immunity is naturally active, significantly limits phage replication, and requires CapV, CdnA, and Cap2 enzyme activities for phage targeting.

**Figure 1.**
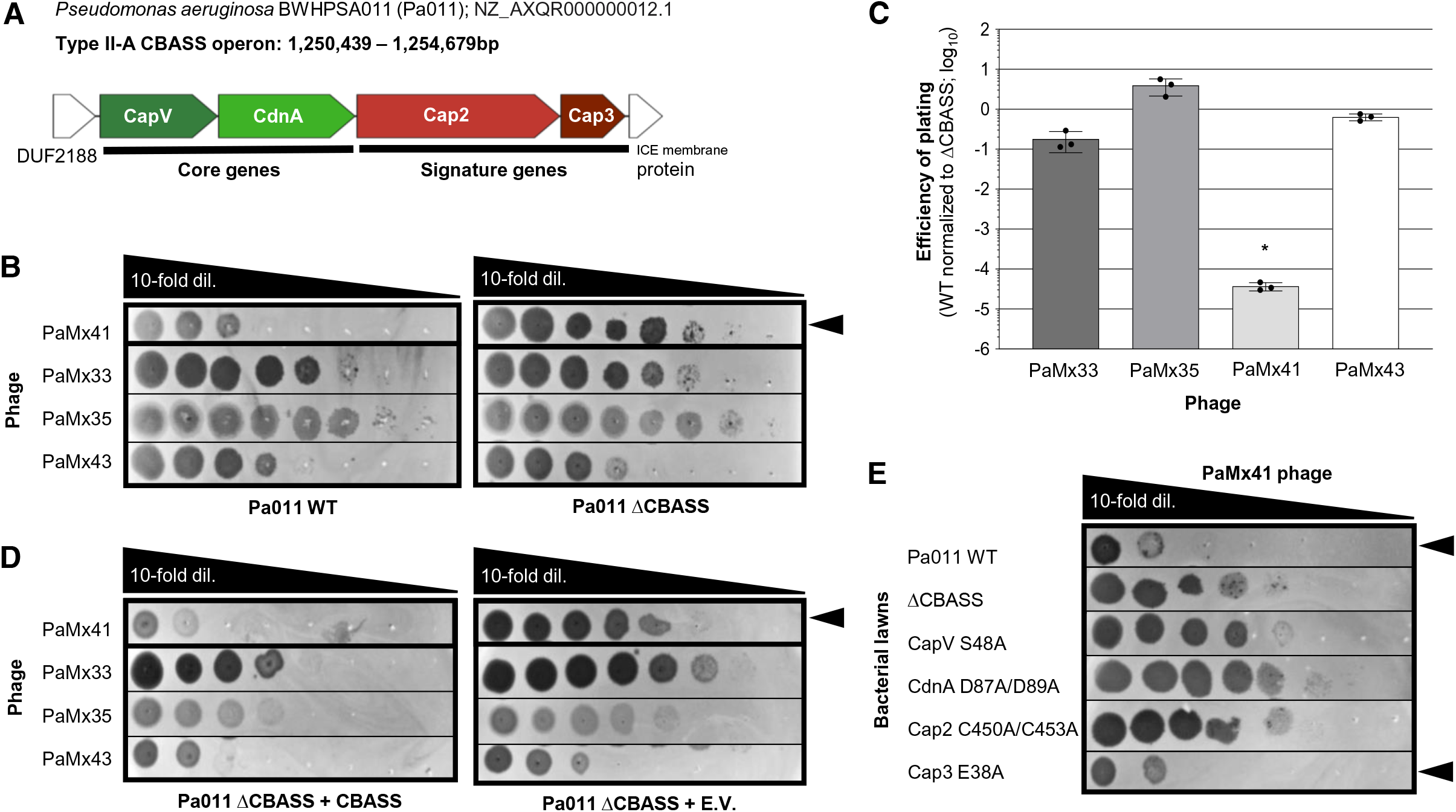
*P. aeruginosa* BWHPSA011 (Pa011) CBASS-based immunity protects against PaMx41 infection. **(A)** Pa011 CBASS operon [CapV (phospholipase), CdnA (cyclase), Cap2 (E1/E2), and Cap3 (JAB)], and adjacent genes [(DUF2188 and ICE (Integrative <u>c</u>onjugative <u>e</u>lement) membrane protein]. **(B)** Plaque assays with the indicated phages spotted in 10-fold serial dilutions on a lawn of Pa011 WT or ΔCBASS; clearings represent phage replication and black arrowhead highlights reduction in PaMx41 WT replication. **(C)** Efficiency of plating was quantified as plaque-forming units (PFU) per ml on Pa011 WT divided by PFU/ml on ΔCBASS (*n*=3). Data are mean ± s.d. (*) Non-parametric ANOVA test yielded a *P* value of <0.0001. **(D)** Plaque assays on a lawn of Pa011 ΔCBASS over-expressing the CBASS operon or empty vector (E.V.). **(E)** Plaque assays on a lawn of Pa011 chromosomal mutants of each CBASS gene. Plaque assays on lawns of Pa011 WT and ΔCBASS were performed in parallel and serve as the controls; black arrowhead highlights reduction in PaMx41 WT replication.

### PaMx41-like phages encode a CBASS antagonist

The phage activator(s) of CBASS immunity is currently unknown. To identify phage genes required for the induction of CBASS immunity, we isolated PaMx41 mutants that escape CBASS. With a frequency of 3.7 × 10^-5^ (Figure 1C), PaMx41 CBASS “escaper” phages were isolated (10 in total) that replicate well on Pa011 WT (Figure 2A). Whole genome sequencing revealed one common mutation in all CBASS escapers, a no-stop extension mutation (X37Q) in *orf24* (Figure 2B). X37Q lengthens gp24 from a 37 amino acid (a.a.) protein to 94 a.a. Interestingly, the CBASS resistant PaMx41-like phages (PaMx33, PaMx35, and PaMx43) naturally encode the 94 a.a. version of gp24 with >98% a.a. identity. While the function of gp24 has not been determined, northern blotting previously revealed that gp24 is expressed in the middle stage of the PaMx41 replication cycle (Cruz-Plancarte et al., 2016).

**Figure 2.**
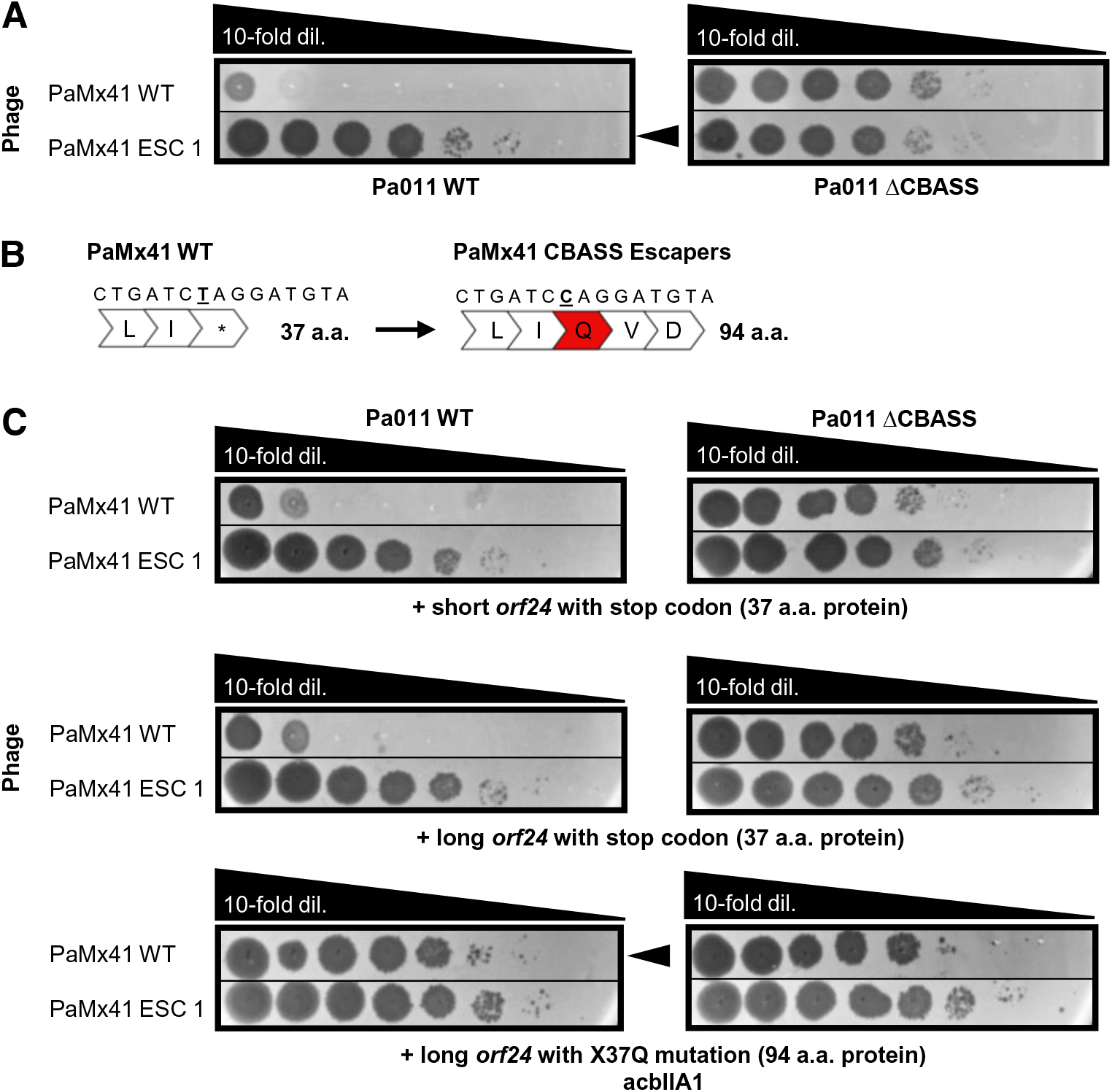
PaMx41 phage mutants escape CBASS immunity. **(A)** Plaque assays with PaMx41 WT phage and an evolved CBASS escaper (ESC) phage spotted in 10-fold serial dilutions on a lawn of Pa011 WT or ΔCBASS; clearings represent phage replication and black arrowhead highlights increase in CBASS escaper phage replication. **(B)** Schematic of the *orf24* no-stop mutation in PaMx41 CBASS escapers phages. The bold underline indicates mutation of thymine (T) to cytosine (C), resulting in a stop codon (TAG, *) to glutamine (CAG, Q) substitution (in red). **(C)** Plaque assays on a lawn of Pa011 WT or ΔCBASS over-expressing the indicated phage genes from the PaMx41 background. Black arrowhead highlights increase in PaMx41 WT phage replication.

To determine whether the short gp24 activates CBASS or the long gp24 inhibits it, constructs expressing the PaMx41 gp24 variants were over-expressed in Pa011 WT and ΔCBASS cells and then plaque assays were performed. The long gp24 increased the titer of the PaMx41 WT phage ≥4 orders of magnitude in the presence of CBASS immunity while the truncated versions (i.e. the short *orf24* or the long *orf24* but with an early stop codon) had no effect (Figure 2C). By contrast, the PaMx41 escaper phage and PaMx33, PaMx35, and PaMx43 phages exhibited high titer on all strains (Figure 2C, S2), demonstrating that the short protein is not a dominant CBASS activator. We next deleted *orf24* from PaMx41, a PaMx41-*orf24^X37Q^* escaper, PaMx33, PaMx35, and PaMx43 using Cas13a genome engineering technology (Guan et al., 2022) (Figure 3A). The titer of the *Δorf24* phages was reduced 2-5 orders of magnitude on Pa011 WT bacterial lawns relative to their plaquing on the ΔCBASS lawn, which could be complemented with the long gp24 expressed *in trans* (Figure 3B). Taken together, these results indicate that the long gp24, or AcbIIA1 (<u>a</u>nti-<u>cb</u>ass type <u>II</u>-<u>A1</u>; PaMx33 NCBI Accession ID: ANA48877) hereafter, inhibits Pa011 CBASS function and is necessary for phage replication in the presence of CBASS immunity.

**Figure 3.**
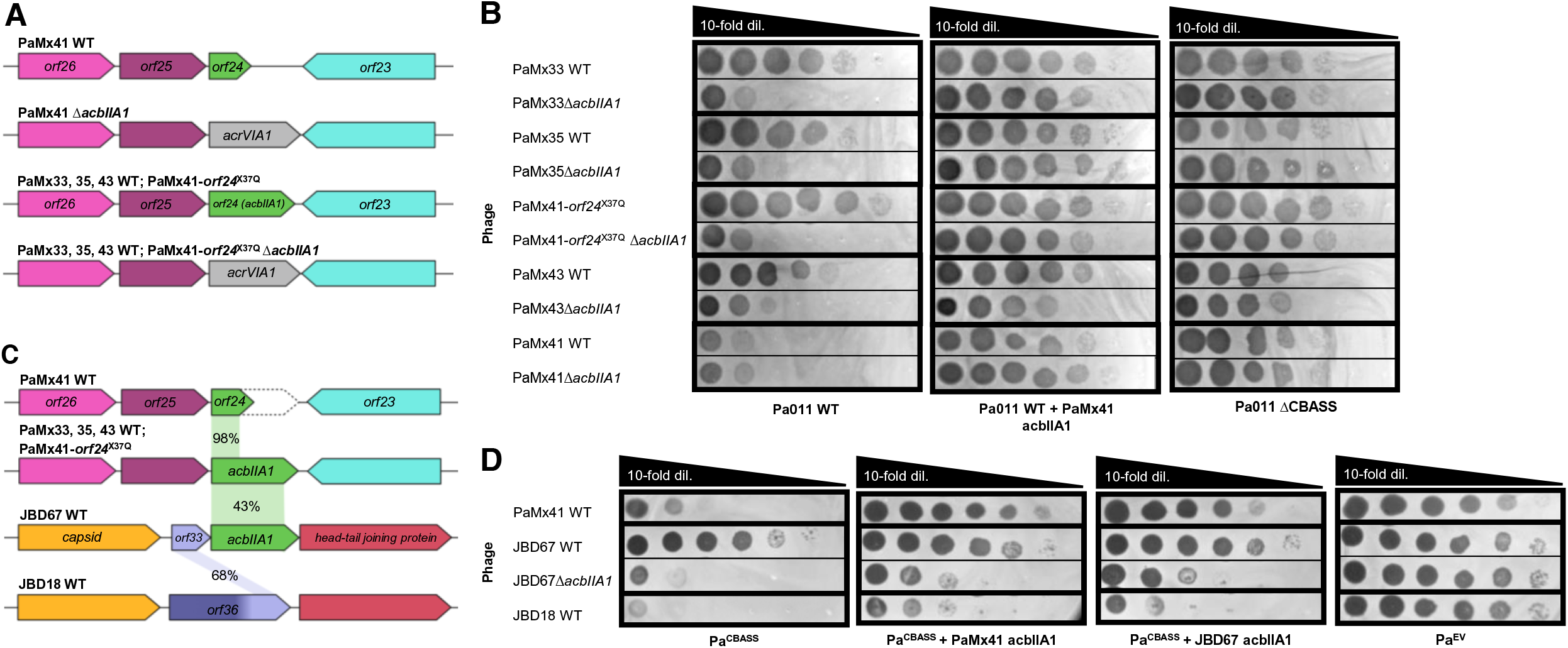
Phage-encoded *acbIIA1* is necessary for replication in the presence of CBASS. **(A)** *acbIIA1* loci in PaMx41-like phage genomes, including WT or *orf24^X37Q^* escaper phages. The Δ*acbIIA1* phages have the *acbIIA1* gene substituted with the type VI-A anti-CRISPR gene (*acrVIA1*) to enable phage engineering (see Methods). **(B)** Plaque assays were performed with the indicated phages spotted in 10-fold serial dilutions on a lawn of Pa011 WT, ΔCBASS, or WT over-expressing PaMx41 *acbIIA1*; clearings represent phage replication. **(C)** Genomic comparison of *acbIIA1* loci in PaMx41-like phages, JBD67, and JBD18 with AcbIIA1 percent amino acid identity shown. Genes with known protein functions are indicated with names, and genes with no predicted function are indicated with “*orf*”. **(D)** Plaque assays with indicated phages spotted in 10-fold serial dilutions on a lawn of *P. aeruginosa* cells (PAO1) with a chromosomally integrated Pa011 CBASS operon (Pa^CBASS^),empty vector (Pa^EV^), or Pa^CBASS^ over-expressing PaMx41 *acbIIA1* or JBD67 *acbIIA1*; clearings represent phage replication.

### acbIIA1 has conserved function in a broadly distributed temperate phage family

Homology searches with AcbIIA1 revealed that it is encoded in a striking number of tailed phages, including those infecting *Pseudomonas, Vibrio, Acinetobacter, Salmonella, Serratia, Erwinia,* and *Escherichia* sp. (including *E. coli* phages T2 and T4, as gene *vs.4*), among others (Figure S3A). The protein has no identifiable domains or predicted molecular functions using BLASTp and hhPred, or AlphaFold2 and DALI servers. A multi-sequence alignment with diverse homologs revealed highly conserved N- and C-termini, with a middle region of varied length and sequence (Figure S3B). *acbIIA1* is commonly encoded by *P. aeruginosa* B3-like temperate siphophages, including JBD67 (with 46% overall a.a. identity, but 75% identity and 95% similarity when analyzing just the N- and C-terminal 55 residues; Figure S3B). Notably, JBD67 is completely unrelated to the PaMx41-like lytic podophages, sharing no other homologous genes. Since JBD67 does not replicate on Pa011 WT or ΔCBASS strains, we integrated the CBASS operon with its native promoter into the chromosome of a *P. aeruginosa* strain (PAO1) that is sensitive to this phage and naturally lacks CBASS. This engineered strain (Pa^CBASS^) reduced PaMx41 WT titer by 3-4 orders of magnitude, and minimally reduced the titers of CBASS resistant phages (PaMx33, PaMx35, PaMx43, or PaMx41-*orf24*^X37Q^ mutant escaper) by 0.5-1 order of magnitude (Figure S4). This modest targeting of these phages was also observed in the native Pa011 strain (Figure 1B-C). JBD67 exhibited resistance to CBASS, while a related phage that naturally lacks the *acbIIA1* gene, JBD18 (Figure 3C), was robustly inhibited (Figure 3D). Deletion of the *acbIIA1* homolog in JBD67 using a modified helicase Cascade-Cas3 system (see Methods, Table S1) sensitized it to CBASS immunity. The JBD67Δ*acbIIA1* and JBD18 phages exhibited ≥5 orders of magnitude reduction in titer in the presence of CBASS, which could be partially complemented *in trans* by *acbIIA1* derived from either JBD67 or PaMx41-*orf24*^X37Q^ (Figure 3D). In parallel, the CBASS sensitive phage PaMx41 WT exhibited full complementation *in trans* by *acbIIA1* derived from either JBD67 or PaMx41-*orf24*^X37Q^ (Figure 3D). These results collectively demonstrate that *acbIIA1* retains its CBASS antagonist function across distinct phage families.

### Phages escape CBASS via mutations in the major capsid gene

Given that the phage activator of CBASS is still unknown, we again attempted to identify phage escapers that lose the CBASS activating factor, or “PAMP” (akin to pathogen-associated molecular patterns in eukaryotes). Using a phage with *acbIIA1* completely removed (PaMx41*ΔacbIIA1*), we searched for escaper plaques on the native Pa011 strain. Phages were first exposed directly to CBASS by plating on Pa011, which yielded no escaper plaques. However, the invisible phage population on the plate was collected, amplified in a ΔCBASS strain and ultimately phage escapers were isolated under CBASS selection (Figure 4A). Whole genome sequencing revealed that every escaper phage (7 in total) had one of four different missense point mutations in the major capsid gene of PaMx41 (*orf11*, NCBI Accession ID: YP_010088887.1; Figure 4B, Table S2). Each escaper phage also had one mutation in a gene of unknown function (*orf52;* Table S3), but no others. The tight clustering of the *orf52* mutations and the silent nature of two of them led us to identify this region as a Type I Restriction-Modification (R-M) site “GT<u>A</u>NNNNNN<u>GTC</u>Y” (positions of mutations underlined, site identical to *P. aeruginosa* enzyme Pae12951I, per REBASE). We surmise that escape of the Type I R-M system resident in Pa011 was also necessary due to the phage being passaged on an alternate host during *acbIIA1* deletion (see Methods). To confirm this hypothesis, we modified the protocol to first passage the PaMx41*ΔacbIIA1* phage in the absence of CBASS immunity to enable host adaptation. CBASS escaper phages were then selected in the presence of CBASS from a high titer population of host adapted phages. Sanger sequencing of the capsid indeed revealed two of the same capsid mutations while mutations in *orf52* no longer appeared (Table S4). To confirm the causality of the capsid mutations for CBASS escape, we used plasmid-based recombination to introduce *de novo* mutations I121A, I121T, and S330P into a naive PaMx41Δ*acbIIA1* phage and confirmed that these mutations induced CBASS escape (Figure 4C-D). However, the over-expression of the PaMx41 WT major capsid protein *in trans* did not induce CBASS targeting of the escaper phages, nor did over-expression of the PaMx41 mutant major capsid protein induce escape during infection with WT phage (data not shown).

**Figure 4.**
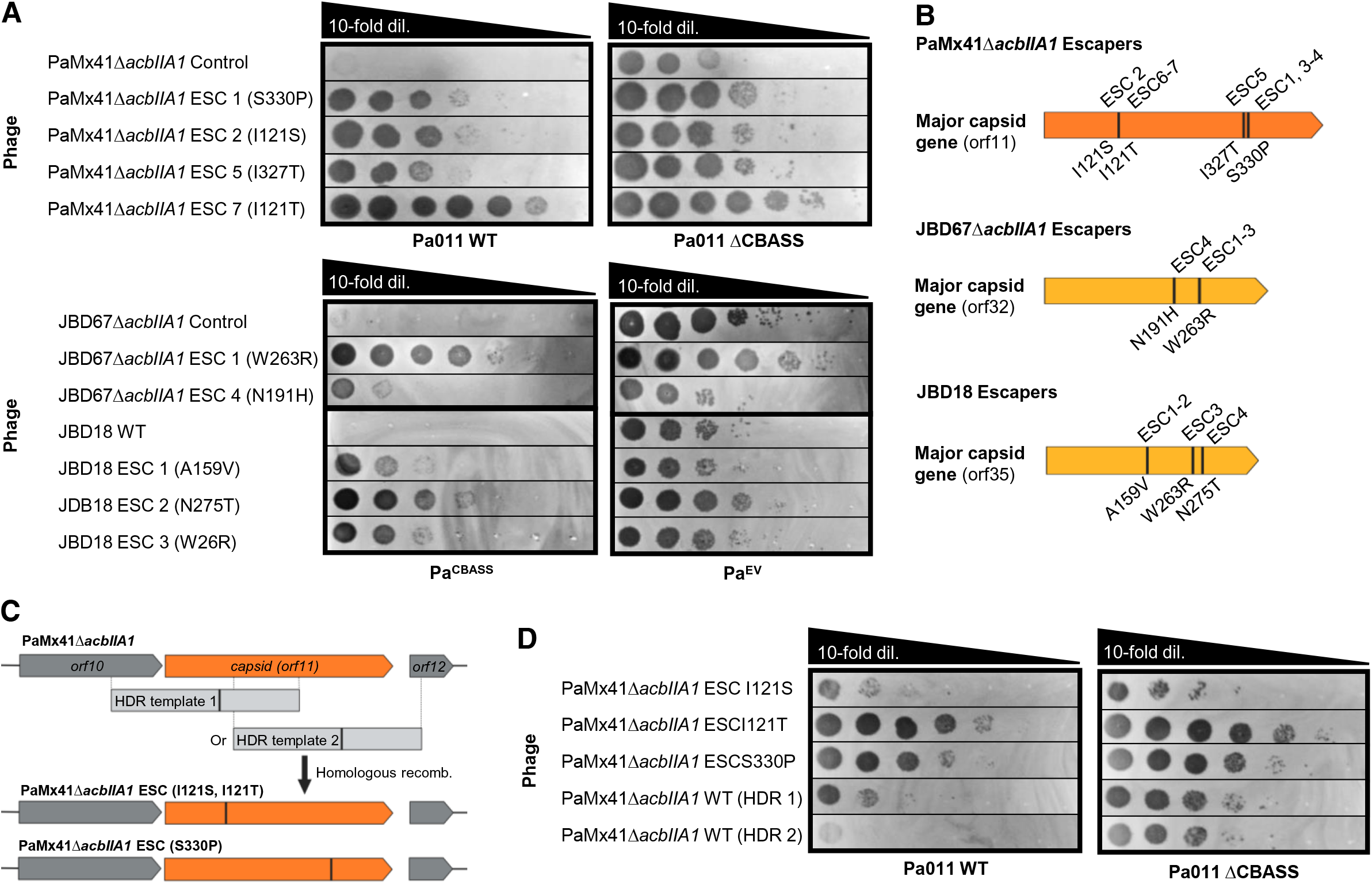
CBASS escaper phages have mutations in their major capsid genes. **(A)** Plaque assays were performed with the indicated control/WT and escaper (ESC) phages spotted in 10-fold serial dilutions on lawns of bacteria expressing CBASS (left) or lacking CBASS (right); clearings represent phage replication. **(B)** Schematic of capsid genes with corresponding missense mutations and associated CBASS escaper phages. **(C)** Schematic of *in vivo* homologous recombination of parental phages with homology-directed repair (HDR) template 1 (encoding I121S or I121T capsid mutations), or template 2 (S330P capsid mutation), and resultant engineered/recombinant phages. **(D)** Plaque assays with recombinant phages possessing major capsid mutations, or WT capsid controls, spotted on lawns of Pa011 WT or ΔCBASS.

CBASS escapers were also isolated from phages JBD67Δ*acbIIA1* and JBD18 using the Pa^CBASS^ strain (Figure 4A), and then the phages were whole genome sequenced. Strikingly, missense mutations in the genes encoding the major capsid protein [*orf32* in JBD67 (NCBI Accession ID: YP_009625956), and *orf35* in JBD18 (NCBI Accession ID: AFR52188); Figure 4B, Table S2] were the only mutations in these CBASS escapers. Given that the major capsid protein of the PaMx41 phage shares no significant amino acid identity with the major capsid protein from JBD67 and JBD18, yet the missense mutations all converge here, it is likely that the major capsid protein is a CBASS activator or target. Follow-up studies are needed to determine the mechanistic connection between the major capsid proteins, the isolated mutants, and CBASS immunity.

## Discussion

The discovery and characterization of CBASS in bacteria marked an exciting connection between prokaryotic and eukaryotic immunity (Cohen et al., 2019; Whiteley et al., 2019). However, many mechanistic and regulatory questions remain in this nascent field. Our identification of endogenous CBASS immunity in *P. aeruginosa*, and the widespread phage gene *acbIIA1* required for phage replication in its presence, provides direct evidence of CBASS as a bona fide anti-phage bacterial immune system. *acbIIA1* is expressed at the middle stage of the PaMx41 phage replication cycle (Cruz-Plancarte et al., 2016), yet the molecular mechanism of CBASS antagonism by AcbIIA1 is currently unclear and warrants further investigation. Furthermore, we observed that *acbIIA1* is encoded adjacent to the capsid gene in phage JBD67, yet its genomic position is highly variable across many of the other phages that encode the gene. This finding demonstrates that there is not an obligate genomic association with the putative CBASS activator.

Upon deletion of *acbIIA1* from the genomes of PaMx41 or JBD67, these phages succumb to CBASS immunity, but rare mutants escape CBASS via missense mutations in the major capsid gene. These results suggest that detection – either direct or indirect – of the late-expressed major capsid protein activates the CdnA enzyme, which is consistent with previous data showing cGAMP levels increasing 40 minutes after P1 phage infection in *E. coli* (Cohen et al., 2019). Moreover, these mutations mirror the recent identification of major capsid mutations in phage T5, which enables evasion of the Pycsar cCMP-based anti-phage bacterial immune system (Tal et al., 2021). The aforementioned study also noted that over-expression of the T5 WT major capsid protein did not induce Pyscar-specific toxicity (Tal et al., 2020), which was similarly observed in our CBASS model system. Therefore, while the mutant phage data suggest that CBASS and Pycsar likely serve as “back-up” immune systems when front-line defenses have failed, the molecular mechanism of activation remain unclear. This interestingly stands in contrast to the early detection of dsDNA during viral infection of eukaryotic cells via the cGAS-STING signaling pathway (Ishikawa et al., 2008; Burdette et al., 2011; Li et al., 2013; Wu et al., 2013). However, these findings don’t rule out the possibility that CBASS blocks biogenesis or assembly of the capsid protein and that our isolated phage mutants escape that CBASS targeting activity. Follow-up studies are needed to define the role of the phage capsid within the context of CBASS immunity.

The human pathogen *Pseudomonas aeruginosa* is an attractive host for the study of endogenous phage-CBASS interactions and evolution, and will likely prove useful in the future for understanding how all types of CBASS are regulated. Moreover, the identification of AcbIIA1 and other anti-CBASS genes in the future may fortify phage therapeutics when used against this and other bacterial pathogens that rely on CBASS for protection. Meanwhile, understanding the molecular mechanisms that underlie the CBASS phenomena (i.e. inhibition, activation, and escape) in this study will provide vital insights into phage-host co-evolution and the ancestor of eukaryotic cGAS-based anti-viral immunity.

## Acknowledgments

We thank current and past members of the Bondy-Denomy Lab for thoughtful discussions, including Drs. Bálint Csörgo, Lina León, and Yuping Li for their technical expertise and knowledge of bacterial and molecular genetics. Drs. Shweta Karambelkar and Lina León for the CRISPR-Cas3 helicase attenuated technology to delete JBD67 *acbIIA1*, and Matt Johnson for bioinformatic comparison of phage genomes. We also thank Dr. Deborah Hung at the Broad Institute of MIT and Harvard for the sample of the *Pseudomonas aeruginosa* strain BWHPSA011, and Dr. Gabriel Guarneros Pena at Centro de Investigacion y de Estudios Avanzados for the PaMx33, PaMx35, PaMx41, and PaMx43 phages. Schematics for figures were created on BioRender.com.

## Author contributions

E.H. conceived the project, designed and performed all experiments, and wrote the manuscript. J.A. contributed to cloning experiments, J.G. engineered PaMx41 with Cas13a. S.S. and H.C. contributed to execution and analysis of the whole genome sequencing data. J.B.-D. supervised the project, designed experiments, and wrote the manuscript.

## Methods

### Identification of CBASS operons

tBLASTn was used to query the amino acid sequence of eight known CD-NTases (CdnA-H) against sequenced *Pseudomonas aeruginosa* genomes contained in the NCBI and IMG databases as well as our sequenced UCSF clinical isolates. Proteins with >24.5% amino acid sequence identity to a validated CD-NTase were accepted as “hits” (Whiteley *et al*., 2019), leading to the identification of >300 CBASS operons in 252 distinct *P. aeruginosa* strains. The *P. aeruginosa* BWHPSA011 (Pa011) strain contains a Type II-A CBASS operon in contig 12 (NCBI Genome ID: NZ_AXQR000000012.1) ranging from 1250439-1254679bp, with CapV (phospholipase effector, NCBI Gene ID: Q024_30602), CdnA (cyclase, Q024_30601), Cap2 (E1/E2, Q024_30600), and Cap3 (JAB, intergenic region 1250436-1250912bp).

### Bacterial strains and growth conditions

The strains, phages, plasmids, and guides used in this study are listed in the Supplementary Table 2. The *P. aeruginosa* strains (BWHPSA011 and PAO1) were grown in LB at 43°C and the *E. coli* strains (DH5ɑ and SM10) were grown in LB at 37°C both with aeration at 225 rpm. Plating was performed on LB solid agar with 10 mM MgSO_4_, and when indicated, gentamicin (50 µg ml^-1^ for *P. aeruginosa* and 15 µg ml^-1^ for *E. coli*) was used to maintain the pHERD30T plasmid. Gene expression was induced by the addition of L-arabinose (0.01% final for BWHPSA011 bacterial genes and 0.1% for phage genes, unless otherwise specified).

### Episomal gene expression

The shuttle vector that replicates in *P. aeruginosa* and *E. coli*, pHERD30T (Qui et al., 2008) was used for cloning and episomal expression of genes in *P. aeruginosa* BWHPSA011 (Pa011) or PAO1 strains. This vector has an arabinose-inducible promoter and a selectable gentamicin marker. For large genes or operons, the vector was digested with SacI and PstI restriction enzymes and purified. Inserts were amplified by PCR using bacterial overnight culture or phage lysate as the DNA template, and joined into the pHERD30T vector at the SacI-PstI site by Gibson Assembly (NEB) following the manufacturer’s protocol. For small genes, the pHERD30T vector was digested with NcoI and KpnI restriction enzymes and purified. Inserts were amplified by PCR using bacterial overnight culture or phage lysate as the DNA template. The inserts were subjected to digestion with NcoI and KpnI, purified, and then joined into the pHERD30T vector at the NcoI-KpnI site by ligation (NEB) following the manufacturer’s protocol. The resulting plasmids were used to transform *E. coli* DH5ɑ. All plasmid constructs were verified by sequencing using primers that annealed to sites outside the multiple cloning site. *P. aeruginosa* cells were electroporated with the pHERD30T constructs and selected on gentamicin.

### Chromosomal CBASS integration

For chromosomal insertion of the Pa011 CBASS operon, the integrating vector pUC18-mini-Tn7T-LAC (Choi *K.-H. et al.*, 2006) and the transposase expressing helper plasmid pTNS3 (Choi *K.-H. et al.* 2008) were used to insert CBASS at the Tn7 locus in *P. aeruginosa* PAO1 strain. The vector was linearized using around-the-world PCR, treated with DpnI, and then purified. Two overlapping inserts encompassing the CBASS operon were amplified by PCR using Pa011 overnight culture as the DNA template, and joined into the pUC18-mini-Tn7T-LAC vector a the SacI-PstI restriction enzyme cut sites by Gibson Assembly (NEB) following the manufacturer’s protocol. The resulting plasmids were used to transform *E. coli* DH5ɑ. All plasmid constructs were verified by sequencing using primers that annealed to sites outside the multiple cloning site. *P. aeruginosa* PAO1 cells were electroporated with pUC18-mini-Tn7T-LAC and pTNS3 and selected for on gentamicin. Potential integrants were screened by colony PCR with primers PTn7R and PglmS-down, and then verified by sequencing using primers that anneal to sites outside the attTn7 site. Electrocompetent cell preparations, transformations, integrations, selections, plasmid curing, and FLP-recombinase-mediated marker excision with pFLP were performed as described previously (Choi K.-H. et al., 2006).

### Chromosomal mutants of *P. aeruginosa* BWHPSA011

The allelic exchange vector that replicates in *P. aeruginosa* and *E. coli*, pMQ30 (Shanks R.M.Q. et al., 2*0*06) was used for generating the chromosomal CBASS knockout and CBASS mutant genes in *P. aeruginosa* BWHPSA011 (Pa011). Vector was digested with HindIII and BamHI restriction enzymes and purified. For the CBASS knockout strain, homology arms >500bp up- and downstream of CBASS operon were amplified by PCR using Pa011 overnight culture as the template DNA. For the CBASS gene mutant strains, homology arms >500bp up- and downstream of CBASS gene catalytic residue(s), with the appropriate mutant nucleotides, were amplified by PCR using Pa011 overnight culture as the template DNA. Previously identified catalytic residues in *Escherichia coli* TW11681 (NZ_AELD01000000) (Cohen *et al.,* 2019) were used to aid the identification of the catalytic residues in *Pseudomonas aeruginosa* BWHPSA011. Multiple Sequence Comparison by Log-Expectation (MUSCLE) and NCBI Multiple Sequence Alignment Viewer were subsequently used to validate conserved catalytic residues between *P. aeruginosa* BWHPSA011, *Vibrio cholerae* El Tor N16961 (NC_002505.1), and *E.coli* TW11681.The inserts were joined into the pMQ30 vector at the HindIII-BamIII restriction enzyme cut sites by Gibson Assembly (NEB) following the manufacturer’s protocol. The resulting plasmids were used to transform *E. coli* DH5ɑ. All plasmid constructs were verified by sequencing using primers that annealed to sites outside the multiple cloning site. *E. coli* SM10 cells were electroporated with pMQ30 constructs and transformants selected for on gentamicin. *E. coli* SM10 harboring the pMQ30 construct were mated with Pa011 to transfer the plasmid and enable allelic exchange. Potential mutant Pa011 strains were subjected to a phenotype cross streak screen with PaMx41-like phages and then verified by sequencing using primers that anneal to sites outside of the homology arms. Electrocompetent cell preparations, transformations, selections, and plasmid curing were performed as described previously (Hmelo L.R. et al., 2015).

### Phage growth

All phages were grown at 37°C with solid LB agar plates containing 20 ml of bottom agar containing 10 mM MgSO_4_ and any necessary inducers or antibiotics. Phages were initially grown on the permissible host *P. aeruginosa* PAO1 WT, which naturally lacks CBASS. 150 µl of overnight cultures of PAO1 were infected with 10 µl of low titer phage lysate (>10^4-7^ pfu/ml) and then mixed with 3 ml of 0.7% top agar 10 mM MgSO_4_ for plating on the LB solid agar. After incubating at 37°C overnight, individual phage plaques were picked from top agar and resuspended in 200 μl SM phage buffer. For high titer lysates, the purified phage was further amplified on LB solid agar plates with PAO1 WT. After incubating 37 °C overnight, SM phage buffer was added until the solid agar lawn was completely covered and then incubated for 5-10 minutes at room temperature. The whole cell lysate was collected and a 1% volume of chloroform was added, and then left to shake gently on an orbital shaker at room temperature for 15 min followed by centrifugation at 10,000 x g for 3 min to remove cell debris. The supernatant phage lysate was stored at 4°C for downstream assays.

### Plaque assays

Plaque assays were conducted at 37°C with solid LB agar plates. 150 µl of overnight bacterial culture was mixed with top agar and plated. Phage lysates were diluted 10-fold then 2 µl spots were applied to the top agar after it had been poured and solidified.

### Isolation of CBASS phage escapers

For identifying PaMx41 WT phage escapers of CBASS, 150 µl of overnight cultures of the *P. aeruginosa* strain BWHPSA011 (Pa011) were infected with 10 µl of high titer phage lysate (>10^9^ pfu/ml) and then plated on LB solid agar. After incubating at 37°C overnight, 10 individual phage plaques were picked from top agar and resuspended in 200 μl SM phage buffer. Phage lysates were purified for three rounds using the CBASS expressing strain. Three PaMx41 WT control phages were picked, purified, and propagated in parallel by infecting the Pa011 *Δ*CBASS strain. To validate the phage identity, PCR and Sanger sequencing were performed on nucleotide sequences unique to PaMx41.

To identify PaMx41 *ΔacbIIA1*, JBD67 *ΔacbIIA1*, and JBD18 WT phage escapers of CBASS, 150 µl of overnight cultures of the CBASS expressing strains (Pa011 WT or Pa^CBASS^) were infected with 10 µl of high titer phage lysate (>10^9^ pfu/ml) and plated on LB solid agar. After incubating at 37°C overnight, no obvious plaques were observed. SM phage buffer was added to the entire lawn and whole cell lysate collected. Next, to propagate the mutant escaper phage population, 150 µl of overnight cultures of the *P. aeruginosa* strains lacking CBASS (Pa011 *Δ*CBASS or Pa^EV^) were infected with 10 µl of the phage lysates and plated on LB solid agar. After incubating at 37°C overnight, SM phage buffer was added to the entire lawn and whole cell lysate collected. Lastly, to isolate individual escaper plaques, 150 µl of overnight cultures of the CBASS expressing strains (Pa011 WT or Pa^CBASS^) were infected with 10 µl of the previously collected phage lysate and plated on LB solid agar. After incubating at 37°C overnight, at least four individual phage plaques were picked from top agar and resuspended in 200 μl SM phage buffer. Phage lysates were purified for three rounds using the CBASS expressing strain. At least two control or WT phages were picked, purified, and propagated in parallel by infecting the Pa011 *Δ*CBASS or Pa^EV^ strains. To validate the phage identity, PCR and Sanger sequencing were performed on nucleotide sequences unique to each phage.

### Whole genome sequencing (WGS) and analysis

Genomic DNA from phage lysates was extracted using a modified SDS/Proteinase K method. 200 μL high titer phage lysate (>10^9^ pfu/ml) was mixed with an equal volume of lysis buffer (10 mM Tris, 10 mM EDTA, 100 μg/mL protease K, 100 μg/mL RNaseA, 0.5% SDS) and incubated at 37°C for 30 min, and then 55°C for 30 min. Preps were further purified using the DNA Clean & Concentrator Kit (Zymo Research). DNA was quantified using the Qubit 4.0 Fluorometer (Life Technologies). 20-100 ng genomic DNA was used to prepare WGS libraries using the Illumina DNA Prep Kit (formerly known as Illumina Nextera Flex Kit) using a modified protocol that utilized 5x reduced quantities of tagmentation reagents per prep, except for the bead washing step with Tagment Wash Buffer (TWB), where the recommended 100 μL of TWB was used. Subsequent on-bead PCR indexing-amplification of tagmented DNA was performed using 2x Phusion Master Mix (NEB) and custom-ordered indexing primers (IDT) matching the sequences from the Illumina Nextera Index Kit. Each 50 μL reaction was split in two tubes, amplified for 9 and 12 cycles respectively. Libraries were further purified by agarose gel electrophoresis; DNA was excised around the ∼400 bp size range and purified using the Zymoclean Gel DNA Recovery Kit (Zymo Research). Libraries were quantified by Qubit and the 9-cycle reaction was used unless the yield was too low for sequencing, in which case the 12-cycle reaction was used. Libraries were pooled in equimolar ratios and sequenced with Illumina MiSeq v3 reagents (150 cycles, Read 1; 8 cycles, Index 1; 8 cycles, Index 2). WGS data were demultiplexed either on-instrument or using a custom demultiplexing Python script (written by Dr. Nimit Jain), and trimmed using cutadapt (v 3.4) to remove Nextera adapters. Trimmed reads were mapped using Bowtie 2.0 (--very-sensitive-local alignments) and alignments were visualized using IGV (v 2.9.4). Variants were detected using the SeqDiff program (https://github.com/hansenlo/SeqDiff).

### CRISPR-Cas13a phage gene editing

Construction of template plasmids for homologous recombination and selection of engineered phages via the CRISPR-Cas13a system were performed as described previously (Guan et al., 2022). Specifically, homology arms of >500bp up- and downstream of PaMx41 *acbIIA1* were amplified by PCR using PaMx41 WT phage genomic DNA as the template. The *acrVIA1* gene was amplified from plasmid pAM383 (Meeske et al., 2020), a gift from Luciano Marraffini, The Rockefeller University. PCR products were purified and assembled as a recombination substrate and then inserted into the NheI site of the pHERD30T vector. The resulting plasmids were electroporated into *P. aeruginosa* PAO1 cells. PAO1 strains carrying the recombination plasmid were grown in LB media supplemented with gentamicin. 150 µl of overnight cultures were infected with 10 µl of high titer phage lysate (>10^9^ pfu/ml; PaMx33 WT, PaMx35 WT, PaMx43 WT, PaMx41-*orf24*^X37Q^ or PaMx41 WT) and then plated on LB solid agar. After incubating at 37°C overnight, SM phage buffer was added to the entire lawn and whole cell lysate collected. The resulting phage lysate containing both WT and recombinant phages were titered on PAO1 strains with a chromosomally integrated Type VI-A CRISPR-Cas13a system, and the most efficiently targeting crRNA guide (specific to *orf11*; guide #5) was used to screen for recombinants. PAO1 strains carrying the Cas13a system and crRNA of choice were grown overnight in LB media supplemented with gentamicin. 150 µl of overnight cultures were infected with 10 µl of low titer phage lysate (10^4-7^ pfu/ml), and then plated onto LB solid agar containing 0.3% arabinose and 1 mM isopropyl β-d-1-thiogalactopyranoside (IPTG). After incubating at 37°C overnight, individual phage plaques were picked from top agar and resuspended in 200 μl SM phage buffer. Phage lysates were purified for three rounds using the Cas13a counter-selection strain (guide #5), and further propagated on a complementary Cas13a counter-selection strain (guide #4), to select against Cas13a escaper phages. To confirm whether the phages were recombinants, PCR was performed with the appropriate pairs of primers amplifying the region outside of the homology arms, an internal region of *acrVIA1*, and *acbIIA1* and then subjected to Sanger Sequencing.

### Homologous recombination-mediated mutation of phage genes

Construction of template plasmids for homologous recombination consisted of homology arms >500bp up- and downstream of the mutation of interest encoded in PaMx41 *orf11*. The homology arms were amplified by PCR using PaMx41Δ*acbIIA1* escapers phage genomic DNA as the template, and PaMx41 WT phage genomic DNA as the control template. Template 1 primers were designed to symmetrically flank the PaMx41 *orf11* mutations I121S and I121T, and template 2 primers were designed to symmetrically flank mutation I327T and S330P. PCR products were purified and assembled as a recombineering substrate and then inserted into the SacI-PstI site of the pHERD30T vector. The resulting plasmids were electroporated into *P. aeruginosa* BWHPSA011 (Pa011) ΔCBASS cells. Pa011 strains carrying the recombination plasmid were grown in LB media supplemented with gentamicin. 150 µl of overnight cultures were infected with 10 µl of high titer phage lysate (>10^9^ pfu/ml; PaMx41Δ*acbIIA1*) and then plated on LB solid agar. After incubating at 37°C overnight, SM phage buffer was added to the entire lawn and whole cell lysate collected. The resulting phage lysate containing both WT and recombinant phages were screened on a lawn of Pa011 WT cells harboring an active CBASS system. Specifically, 150 µl of overnight Pa011 WT cultures were infected with 10 µl of low titer phage lysate (10^4-7^ pfu/ml), and then plated onto LB solid agar. After incubating at 37°C overnight, individual phage plaques were picked from top agar and resuspended in 200 μl SM phage buffer. Phage lysates were purified for three rounds using the Pa011 WT strain. To confirm whether the phages were recombinants, PCR was performed with the appropriate pairs of primers amplifying the region outside of the homology arms and subject to Sanger Sequencing.

### Helicase attenuated Cas3 removal of phage genes

Cas3 (Type I-C)-specific guides targeting JBD67 *acbIIA1* were cloned into a pHERD30T-derived vector containing modified I-C repeats as previously described (Csörgő et al., 2020). The guides were electroporated into *P. aeruginosa* PAO1 strains with a chromosomally integrated Type I-C helicase attenuated Cas3 system. JBD67 WT phage lysate was titered on the PAO1 strains and the efficiently targeting crRNA guide (specific to *acbIIA1;* guide #3) was identified. PAO1 strains carrying the Type I-C CRISPR-Cas system with a helicase attenuated Cas3 enzyme and crRNA targeting phage JBD67 *acbIIA1* were grown overnight in LB media supplemented with gentamicin. 150 µl of overnight cultures were infected with 10 µl of high titer phage lysate (>10^9^ pfu/ml; JBD67) and plated on LB agar plates containing gentamicin, 0.1% arabinose, and 1 mM isopropyl β-d-1-thiogalactopyranoside (IPTG). After incubating at 37°C overnight, SM phage buffer was added to the entire lawn and whole cell lysate collected. The resulting phage lysate containing both WT and *acbIIA1* knockout phages were grown on a complementary Cas3 counter-selection strain (guide #4) to select against Cas3 escaper phages. 150 µl of overnight cultures were infected with 10 µl of low titer phage lysate (10^4-7^ pfu/ml; JBD67 WT) and then plated on LB solid agar containing 0.1% arabinose and 1 mM IPTG. After incubating at 37°C overnight, individual phage plaques were picked from top agar and replica-plated onto LB solid agar with PAO1^EV^ and PAO1^CBASS^ strains. JDB67 plaque sizes that were reduced on the PAO1^CBASS^ plate were identified as potential CBASS sensitive phages. Corresponding plaques on the PAO1^EV^ plate were picked and resuspended in 200 μl SM phage buffer. To determine whether the phages harbored deletions in *acbIIA1*, PCR was performed with the appropriate pairs of primers amplifying a ∼1kb region outside of *acbIIA1*.

### Phylogenetic analysis

Phylogenetic reconstructions were conducted similar to previous work in our lab (Marino et al., 2019). Homologs of AcbIIA1 were acquired through 3 iterations of psiBLASTp search the non-redundant protein database, specifying Caudovirales as the search space. Hits with >70% coverage and an E value <0.0005 were included in the generation of the position specific scoring matrix (PSSM). High confidence homologs (>70% coverage, E value < 0.0005) represented in unique species of bacteria were then aligned using NCBI COBALT (Papadopoulos et al., 2007) using default settings and a phylogeny was generated in Cobalt using the fastest minimum evolution method (Desper et al., 2004) employing a maximum sequence difference of 0.85 and Grishin distance to calculate the tree. The resulting phylogeny was then displayed as a phylogenetic tree using iTOL: Interactive Tree of Life (Letunic et al., 2018).

### Comparative genomics of phage

An in-house program was used to generate phage alignments. GenBank files were obtained for the phage genomes of interest. The program reads the GenBank files and calculates nucleotide similarity using NCBI pairwise BLASTn with default parameters. Protein families are defined by clusters obtained by MMSeqs2 “cluster” mode with coverage parameter “-c 0.8”. The program draws ribbons of high nucleotide identity between the phage genomes with high nucleotide identity (>80%) and protein families are given identical colors.

## Supplemental information titles and legends

**Figure S1.**
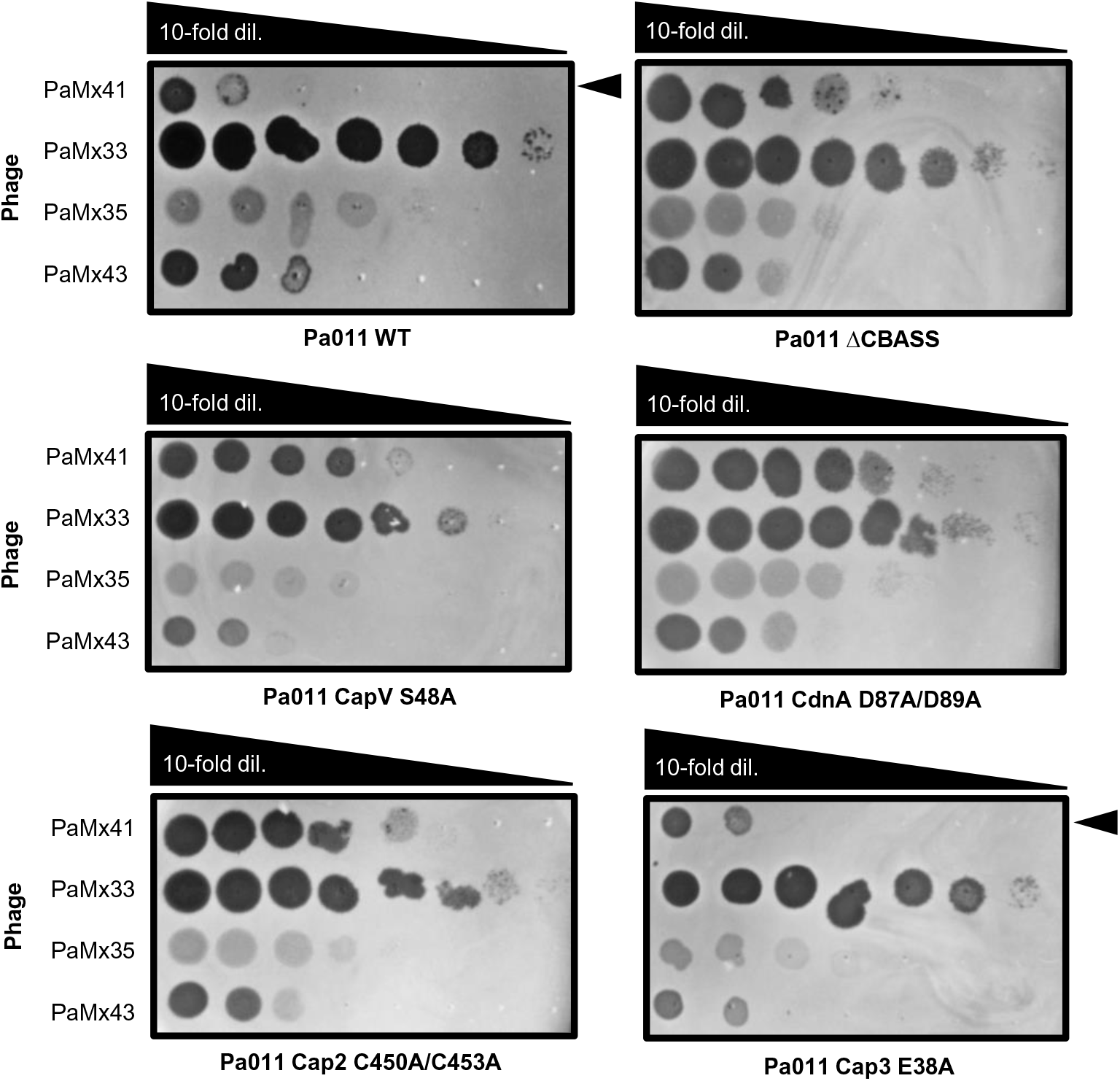
Pa011 CBASS-based immunity provides protection against PaMx41, but not related PaMx33, 35, and 43 phages. Plaque assays were performed with the indicated phages spotted in 10-fold serial dilutions on a lawn of Pa011 chromosomal mutants of each CBASS gene [CapV (phospholipase), CdnA (cyclase), Cap2 (E1/E2), and Cap3 (JAB)]. Plaque assays on lawns of Pa011 WT and ΔCBASS were performed in parallel and served as controls; clearings represent phage replication and black arrowhead highlights reduction in PaMx41 WT replication.

**Figure S2.**
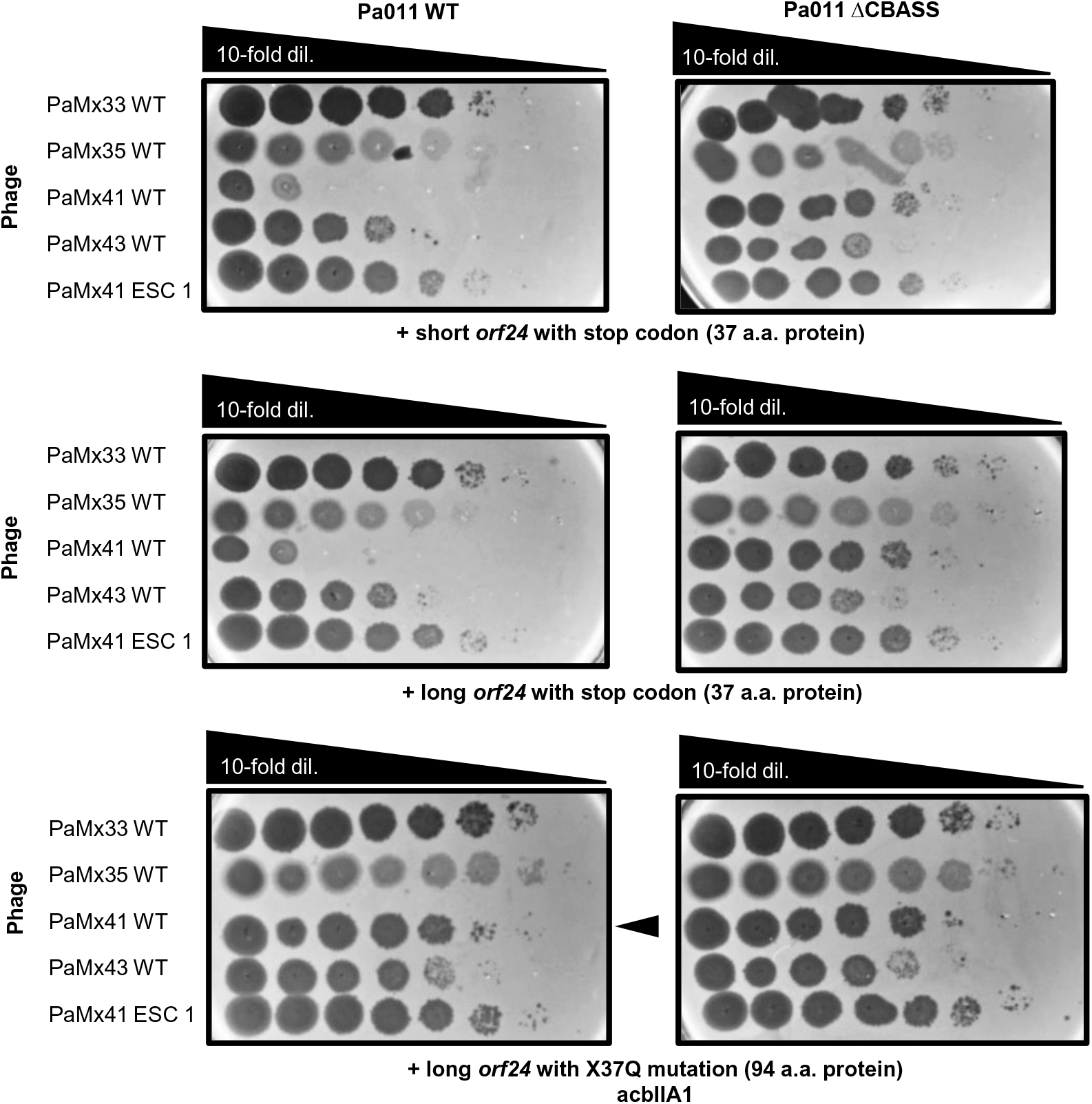
PaMx33, 35, and 43 phages remain resistant to CBASS immunity in the presence of orf24 variants. Plaque assays were performed with PaMx33, 35, 41, and 43 WT phages, as well as an evolved PaMx41 CBASS escaper (ESC) phage, spotted in 10-fold serial dilutions on a lawn of Pa011 WT or ΔCBASS over-expressing the indicated phage genes from the PaMx41 background; clearings represent phage replication and black arrowhead highlights increase in PaMx41 WT replication.

**Figure S3.**
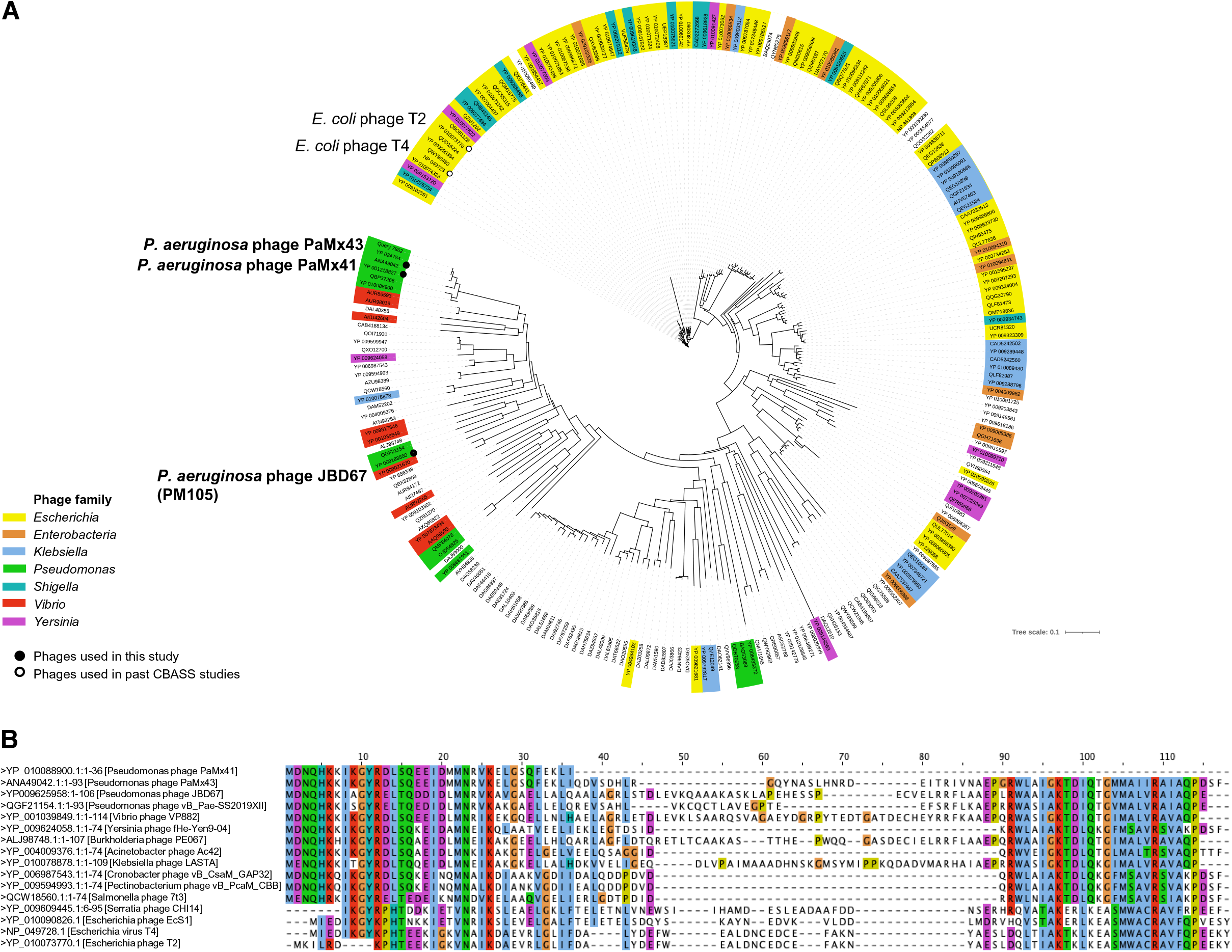
AcbIIA1 is found in a broad diversity of phages. **(A)** Phylogenetic tree of *acbIIA1* across the genomes of 239 tailed phages (*Caudovirales*) following two iterations of PSI-BLAST. Phage families that are colored show some that infect well-known bacterial genera. Phages relevant to this study are bolded. **(B)** Multiple sequence alignment of PaMx41 *orf24*^X37Q^ AcbIIA1, JDB67 AcbIIA1, and other homologous proteins via MAFFT alignment in Jalview. All proteins except T2 and T4 phage homologues were acquired by standard BLASTp, while T2 and T4 were observed on a second round of a PSI-BLAST. Colors indicate an identical amino acid.

**Figure S4.**
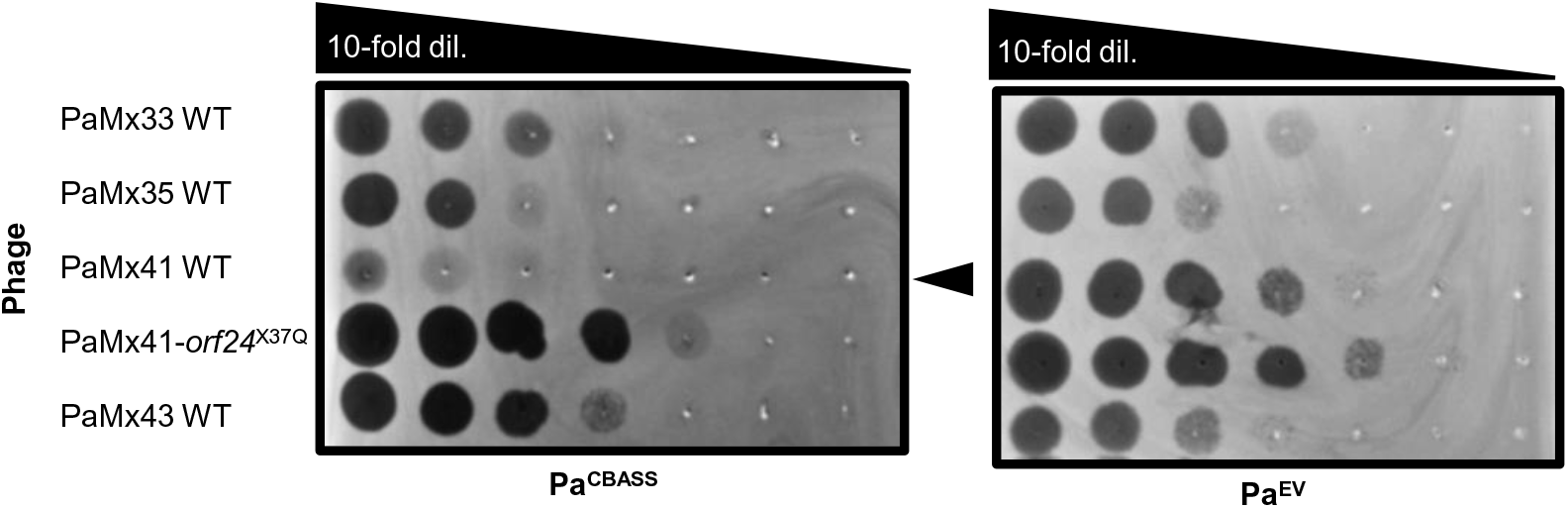
Pa011 CBASS-based immunity is recapitulated in an engineered bacterial strain that chromosomally expresses CBASS. Plaque assays with the indicated phages spotted in 10-fold serial dilutions on a lawn of *P. aeruginosa* cells (PAO1) with a chromosomally integrated Pa011 CBASS operon (Pa^CBASS^) or empty vector (Pa^EV^); clearings represent phage replication and black arrowheadhead highlights reduction in PaMx41 WT replication.

**Table S1.**
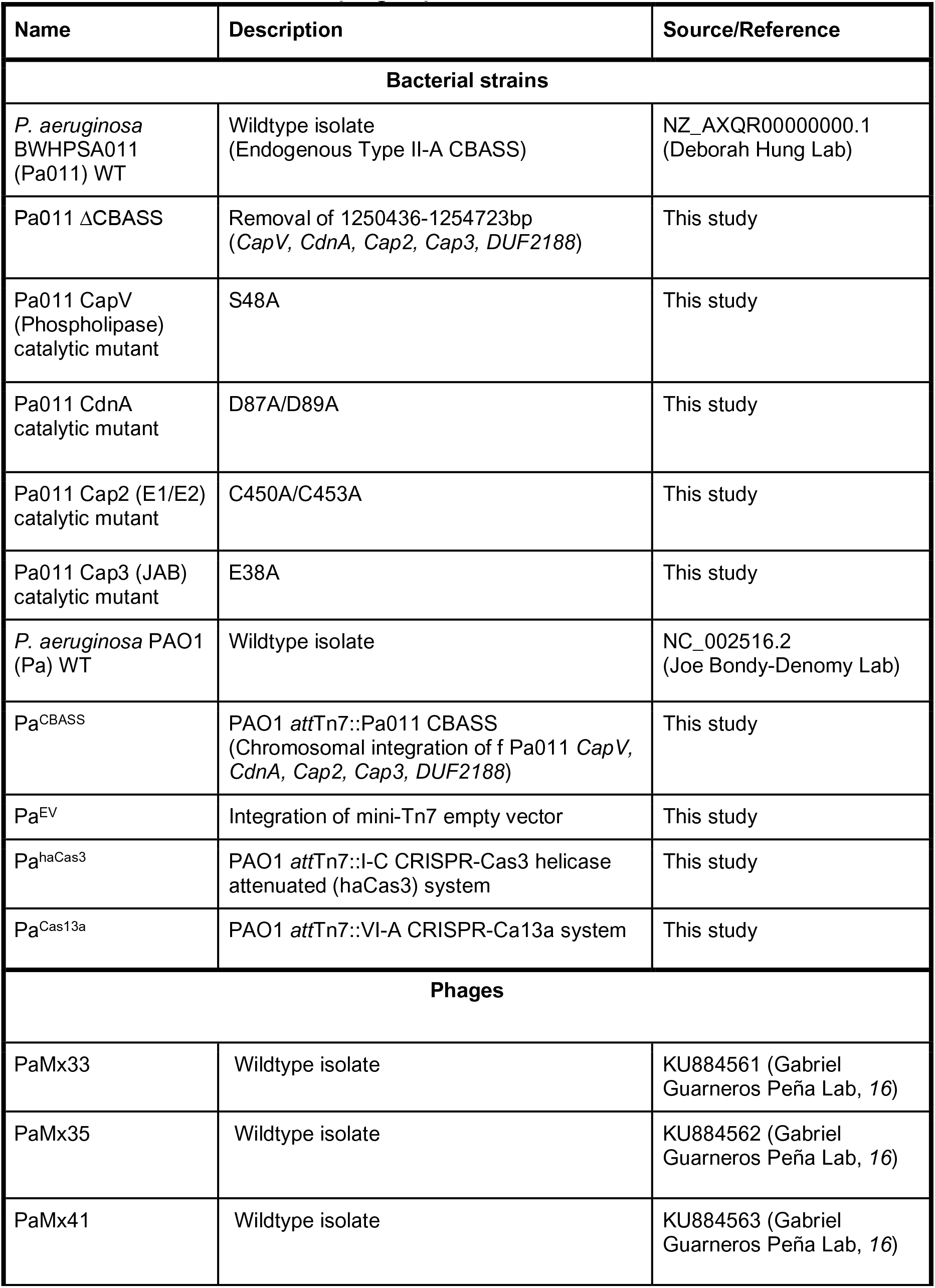
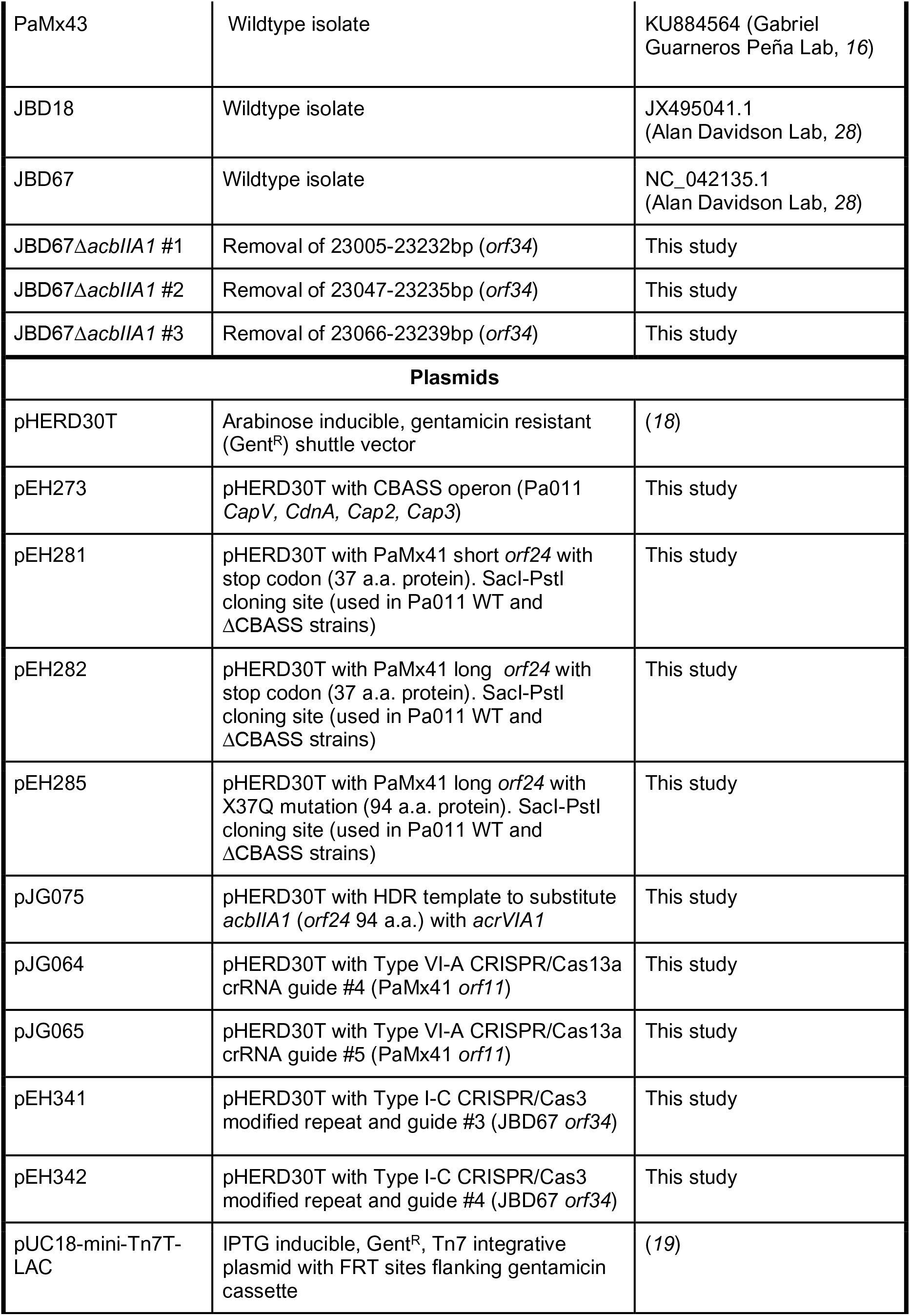
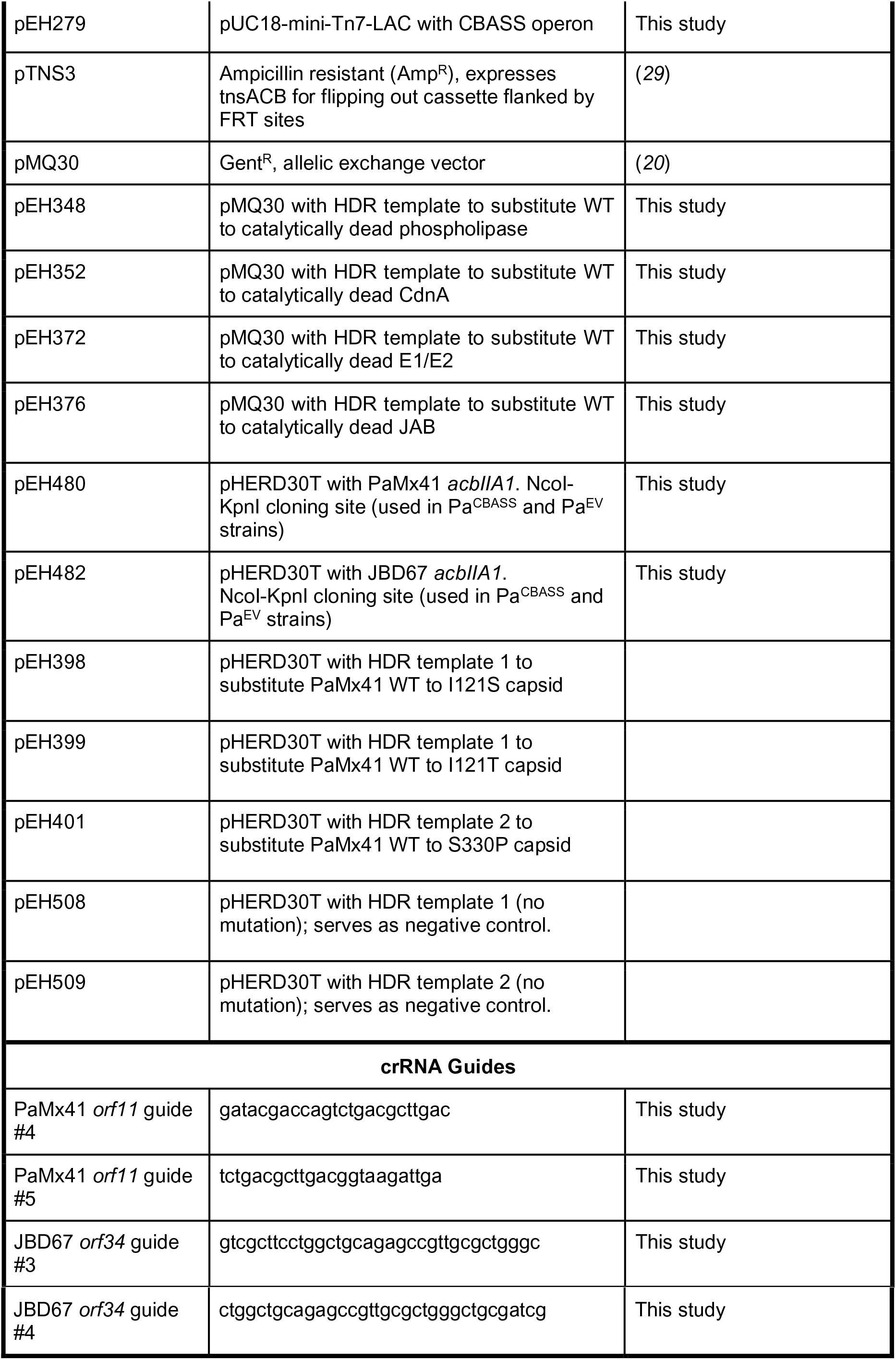
List of bacterial strains, phages, plasmids, and crRNAs.

**Table S2.** List of phage capsid mutations observed via Whole Genome Sequencing.

| Phage name | Nucleotide location | Nucleotide mutation* | Amino acid location | Amino acid mutation |
| --- | --- | --- | --- | --- |
| **PaMx41 Δ*acbIIA1* Escapers (ESC)** |  |  |  |  |
| ESC 1 | 9371 | T→C | 330 | S→P |
| ESC 2 | 8745 | T→G | 121 | I→S |
| ESC 3 | 9371 | T→C | 330 | S→P |
| ESC 4 | 9371 | T→C | 330 | S→P |
| ESC 5 | 9363 | T→C | 327 | I→T |
| ESC 6 | 8745 | T→C | 121 | I→T |
| ESC 7 | 8745 | T→C | 121 | I→T |
| **JBD67 Δ*acbIIA1* ESC** |  |  |  |  |
| ESC 1 | 22601 | T→C | 263 | W→R |
| ESC 2 | 22601 | T→C | 263 | W→R |
| ESC 3 | 22601 | T→C | 263 | W→R |
| ESC 4 | 22385 | A→T | 191 | N→H |
| **JBD18 WT ESC** |  |  |  |  |
| ESC 1 | 22559 | C→T | 159 | A→V |
| ESC 2 | 22559 | C→T | 159 | A→V |
| ESC 3 | 22907 | A→C | 275 | N→T |
| ESC 4 | 22870 | T→C | 263 | W→R |

\*Mutations were recorded if the 100% of the reads detected the change in nucleotide relative to the reference genome.

**Table S3.** List of all mutations in PaMx41 Δ*acbIIA1* escaper (ESC) phage observed via Whole Genome Sequencing.

| Phage name | Nucleotide location | Nucleotide mutation* | Gene name | Amino acid location | Amino acid mutation |
| --- | --- | --- | --- | --- | --- |
| ESC 1 | 9371 | T→C | *orf11* (capsid) | 330 | S→P |
|  | 41715 | G→T | *orf52* (unknown) | 247 | T→I |
| ESC 2 | 8745 | T→G | *orf11* | 121 | I→S |
|  | 41708 | A→G | *orf52* | 249 | A→A |
| ESC 3 | 9371 | T→C | *orf11* | 330 | S→P |
|  | 41715 | G→T | *orf52* | 247 | T→I |
| ESC 4 | 9371 | T→C | *orf11* | 330 | S→P |
|  | 41715 | G→T | *orf52* | 247 | T→I |
| ESC 5 | 9363 | T→C | *orf11* | 327 | I→T |
|  | 41715 | G→T | *orf52* | 247 | T→I |
| ESC 6 | 8745 | T→C | *orf11* | 121 | I→T |
|  | 41716 | T→C | *orf52* | 247 | T→A |
| ESC 7 | 8745 | T→C | *orf11* | 121 | I→T |
|  | 41717 | C→T | *orf52* | 246 | K→K |

\*Mutations were recorded if the 100% of the reads detected the change in nucleotide relative to the reference genome. Note that all control and escaper phages exhibited a single point mutation in *orf51* (C→T mutation at position 41,142; V→I), suggesting that our lab PaMx41 $\Delta acbIIA1$ lineage has a mutation relative to the reference sequence at this site.

**Table S4.** List of all mutations in PaMx41 Δ*acbIIA1* escaper (ESC) phage observed via Sanger Sequencing.

| Phage name | Nucleotide location | Nucleotide mutation | Amino acid location | Amino acid mutation |
| --- | --- | --- | --- | --- |
| ESC 8 | 8745 | T→C | 121 | I→T |
| ESC 9 | 8745 | T→C | 121 | I→T |
| ESC 10 | 8745 | T→C | 121 | I→T |
| ESC 11 | 9363 | T→C | 327 | I→T |
| ESC 12 | 9363 | T→C | 327 | I→T |

